# Detection of intra-tumoral microbiota from transcriptomic sequencing of Asian breast cancer

**DOI:** 10.1101/2024.09.25.614900

**Authors:** Li-Fang Yeo, Audrey Weng Yan Lee, Phoebe Yon Ern Tee, Joyce Seow Fong Chin, Bernard KB Lee, Joanna Lim, Soo-Hwang Teo, Jia Wern Pan

**Affiliations:** Cancer Research Malaysia, Subang Jaya, Malaysia; Department of Mathematical Sciences, Faculty of Science and Technology, Universiti Kebangsaan Malaysia, Selangor, Malaysia

## Abstract

The human microbiome has garnered significant interest in recent years as an important driver of human health and disease. Likewise, it has been suggested that the intra-tumoral microbiome may be associated with specific features of cancer such as tumour progression and metastasis. However, additional research is needed to validate these findings in diverse populations. In this study, we characterized the intra-tumoral microbiota of 883 Malaysian breast cancer patients using transcriptomic data from bulk tumours and investigated their association with clinical variables and immune scores. We found that the tumour microbiome was not associated with breast cancer molecular subtype, cancer stage, tumour grade, or patient age, but was weakly associated with immune scores. We also found that the tumour microbiome was able to predict immune scores in our cohort using random forest models, suggesting the possibility of an interaction between the tumour microbiome and the tumour immune microenvironment in Asian breast cancer.

## 1.0 Introduction

Breast cancer is the most common cancer in women across the majority of countries worldwide. Differences in distribution of genetic (Breast Cancer Association Consortium 2021), lifestyle (Mertens et al. 2023) and reproductive factors (Mao et al. 2023) influence the clinical presentation of breast cancer in different populations. For example, there is a higher prevalence of triple negative breast cancer in women of African descent (Martini et al. 2022), and a higher prevalence of immune enriched breast cancers in women of Asian descent (Pan et al. 2020). Whilst part of these differences may be attributable to differences in population genetics, a large proportion of these differences remain unexplained.

One factor that may potentially explain some of these differences is the microbial community found on and in the human body, also known as the human microbiome. With the advent of next-generation sequencing and decreasing cost to sequence genomes, it has become possible to study the human microbiome in much greater detail. Early studies were mostly focused on characterising the human microbiome (The Human Microbiome Project Consortium 2012), but several recent studies have studied their association with cancer and other diseases. Routy et al. (2018) reported a retrospective cohort study where cancer patients on antibiotics had shorter progression free-survival and overall survival. Restoring gut microbial diversity via live-bacteria supplements (Routy et al. 2018) or faecal microbiota transplant (Riquelme et al. 2019) improved response to anti-PD1 therapy, suggesting that the gut microbiome may play an important role in treatment outcomes.

Researchers have also been interested in the intra-tumoral microbiome and its association with cancer, though this has been more challenging to study due to its low biomass and accessibility. Recently, intratumoral bacteria were found to mostly reside within cancer or immune cells, with each tumour type shown to have a distinct microbiota composition (Nejman et al. 2020). This landmark paper also showed that breast tumours had the richest and most diverse intra-tumoral microbiome, which was associated with clinical subtypes. Other recent studies have demonstrated the ability of intra-tumoral bacteria to induce the migration of cancer cells and promote cancer progression (Galeano Niño et al. 2022) and metastasis (Fu et al. 2022).

Notably, global studies that compare the microbiome of different ethnic groups suggest that population-specific tumour microbiomes may exist. For example, a recent study reported differences between tumoral microbiota composition between Caucasians and African Americans but observed no significant differences for Asians (German et al. 2023). This mirrored findings from Parida et al. (2023), who also found differences between Caucasians and African Americans but not Asians. However, both papers included only a very small number of Asian patients in their analyses. This demonstrates the lack of Asian representation in global cancer microbiome studies, which may in turn lead to false assumptions regarding the generalizability of microbiome studies to the wider Asian population.

In this study, we characterized the tumoral microbiota of 883 Malaysian breast cancer patients using transcriptomic data from bulk tumours and investigated their association with clinical variables and immune scores. We found that the tumour microbiome was not associated with breast cancer molecular subtype, cancer stage, tumour grade, or patient age, but was weakly associated with immune scores. We also found that the tumour microbiome was able to predict immune scores in our cohort using random forest models, suggesting the possibility of interactions between the tumour microbiome and the tumour immune microenvironment in Asian breast cancer.

## 2.0 Methods

### 2.1 Biospecimen collection and data generation

RNA-seq data that was generated by Pan et al. (2020) and Pan et al. (2024) were used to discover the presence of microbes in fresh frozen tumours from 977 breast cancer patients from the Malaysian Breast Cancer (MyBrCa) cohort recruited at Subang Jaya Medical Centre (*n*=843) and University Malaya Specialist Centre (*n*=134), Malaysia. As the sequencing was conducted in two separate batches, the earlier batch was used as a discovery cohort (*n*=558), and the latter as a validation cohort (*n*=419).

### 2.2 Data quality assessment and read alignment

RNA-seq reads that mapped to hs38r42 human genome using STAR aligner were removed (Dobin et al. 2013). Non-human, unmapped reads were retained and mapped to the Kraken2 32GB database (Wood et al. 2019). Relative abundance of microbial reads from Kraken2 were estimated using Bracken (Lu et al. 2017). Read count tables were created for each taxonomic level by using the kreport2mpa.py script from KrakenTools (Lu et al. 2022).

### 2.3 Alpha and beta diversity analyses

Reads were converted to relative abundance. Intra-group (alpha) diversity was determined using the number of observed species and Shannon index. Inter-group (beta) diversity was measured using a Bray-Curtis dissimilarity matrix, plotted using unsupervised, multi-dimensional scaling (MDS) method and visualized on a PCoA (Principal Coordinates Analysis) plot. Shepherd’s stress test was used to measure goodness-of-fit of the model, that is how well the reduced dimensions reflect the original dissimilarity structure. Beta diversity was also measured using supervised ordination, dbRDA (distance-based Redundancy Analysis). PERMANOVA (Permutational Multivariate ANOVA) was used to calculate the differences between groups controlled by covariates. Covariates included in the analysis were PAM50 subtype, age at diagnosis, cancer stage, tumour grade, treatment used, and ethnicity. PERMDISP (Permutational Multivariate Analysis of Dispersion) was calculated to ensure homogenous dispersion was observed in the model as skewed dispersion may confound findings from PERMANOVA.

### 2.4 Differential abundance analyses

Microbial counts at the genus level were filtered for at least 10% prevalence and centre log-ratio (clr) transformed as recommended by Nearing et al. (2022). Differential abundance analyses were done using the compositional data analysis method, namely with ALDEx2, ANCOM-BC2, MaAsLin2, LinDA and Zicoseq. Bacterial taxa were deemed to be significantly different if the FDR-corrected *p*-value was <0.05 in more than two algorithms.

### 2.5 Random Forest modelling

Supervised machine learning using random forest models were tested on a total of 883 samples after filtering out samples with missing data or failed filtering QC. The R packages ‘caret’ (v. 6.0-94) and ‘mikropml’ (v. 1.6.1) were used to train and test random forest models using an 80:20 training:testing split with 5-fold cross-validation repeated five times (the “xgbTree” method was used with the “repeatedCV” option set to 5 repetitions). The R packages ‘mikropml’ (v. 1.6.1) and ‘MLeval’ (v. 0.3) were used to calculate F1, AUROC, precision, sensitivity, specificity and generate plots.

### 2.6 *In vitro* validation of microbial counts

The qRT-PCR assays were performed with QuantiNova SYBR Green RT-PCR Kit (Qiagen) using the Applied Biosystems Real-Time PCR System. Twenty nanograms per microliter (ng/µL) of DNA extracted from tumour samples were used as the template for amplification of the V3V4 region using the forward primer (5’-TCGTCGGCAGCGTCAGATGTGTATAAGAGACAGCCTACGGGNGGCWGCAG-3’) and reverse primer (5’-GTCTCGTGGGCTCGGAGATGTGTATAAGAGACAGGACTACHVGGGTATCTAATCC-3’) sequences obtained from Klindworth et al. (2013). Reaction mixtures consisted of 10µL master mix, 3µL each of forward and reverse primers, and 4µL of DNA template. *Escherichia coli* gDNA was employed as a positive control, OKF6/TERT1 gDNA as a negative control, and water was used as a blank control. Cycles consisted of the following regime: 2 min at 50°C, 10 min at 95°C, 40 cycles of 15 s at 95°C and 30 s at 60°C, followed by 15 s at 95°C, 1 min at 60°C, 30 s at 95°C, and 15 s at 60°C for melt curve analysis. A total of 5µL of the final qRT-PCR amplicons were subjected to agarose gel electrophoresis in a 2% gel at 100 volts for 30 minutes and visualised under ultraviolet (UV) transillumination on the Azure Biosystems Imaging System. *E. coli* gDNA was serially diluted and ran in triplicates on the qRT-PCR system. The measured threshold cycle (Ct) values were plotted against calculated copy numbers for each reaction. Ct values from the V3V4 qRT-PCR analyses were used to estimate copy numbers of total bacteria present in tumour samples based on the standard curve. Estimated copy numbers were then compared with microbial counts of corresponding samples and used to generate a correlation plot.

## 3.0 Results

### 3.1 Alpha, beta diversity and most prevalent genera in the Malaysian breast tumour microbiome

Using our discovery cohort (*n*=558), bacteria read counts were converted to relative abundance to observe the overall distribution of each taxonomy when grouped by PAM50 subtype (Figure 1A). Proteobacteria, Firmicutes, Actinobacteria, and Heunggongvirae were the most dominant phyla of the breast tumour microbiota. There was significant heterogeneity, where some phyla, such as Acidobacteria, were observed in some samples but were completely absent in others. The top ten most abundant genera in our discovery cohort by median read counts were *Pseudomonas, Siphoviridae, Bacillus, Escherichia, Klebsiella, Streptomyces, Priestia, Cutibacterium, Serratia*, and *Acinetobacter*.

**Figure 1.**
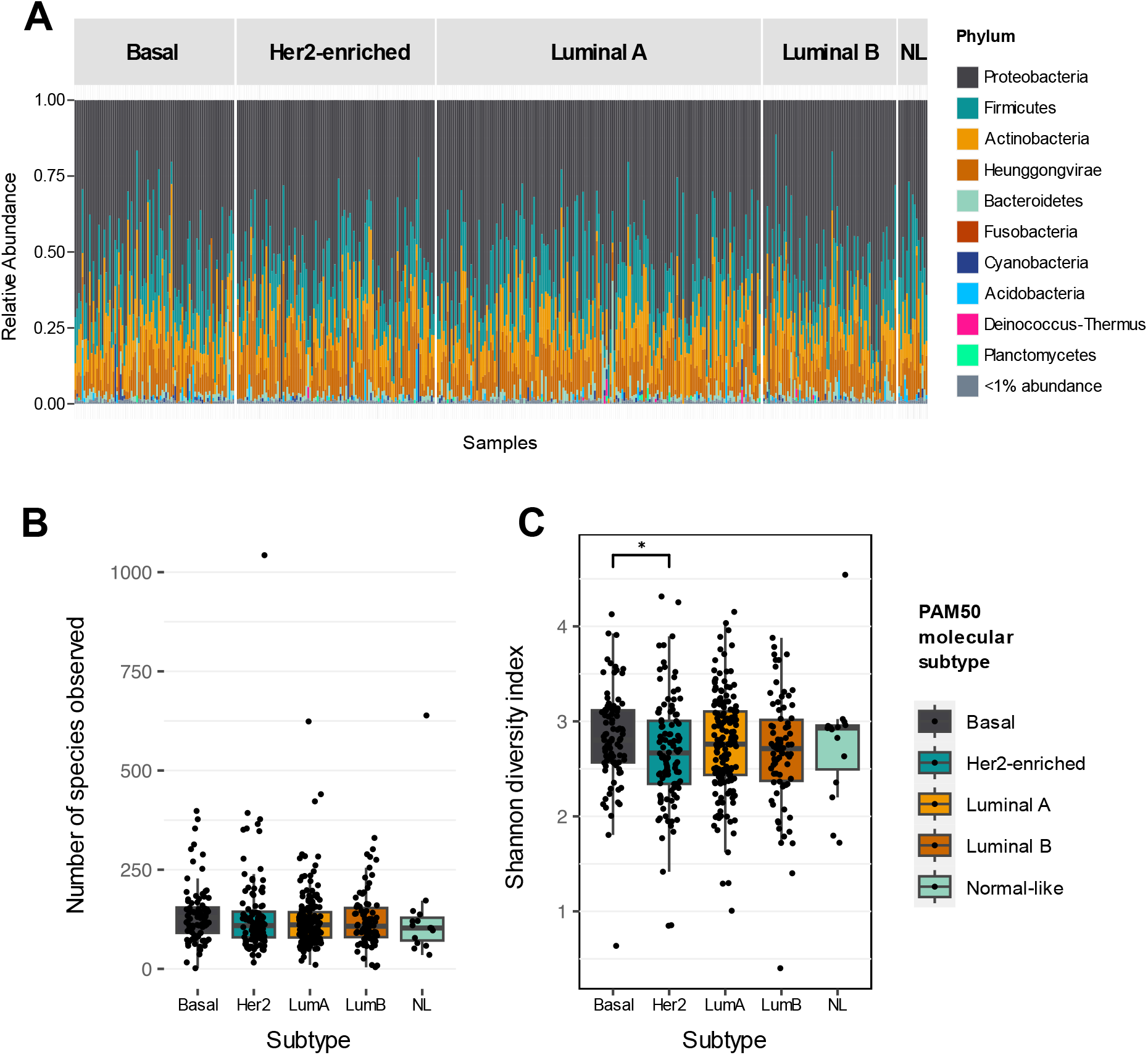
Detection of intratumoral microbiota from transcriptomic sequencing of Asian breast cancer samples. **(A)** Relative abundance of microbial phyla detected in breast cancer samples from the MyBrCa discovery cohort (n=558), grouped by PAM50 molecular subtype (NL = normal-like). Also shown are the number of species observed **(B)** and the Shannon diversity index (**C)** for detected intratumoral microbiota across different breast cancer molecular subtypes.

The intra-group diversity of breast tumour microbiota was mostly homogenous when comparing between molecular subtypes. A slightly higher diversity was observed in Basal subtype when using observed number of species as alpha diversity metric, although it did not reach statistical significance (Figure 1B, *p*>0.05). The diversity of the Basal subtype microbiota was significantly higher than the Her2 subtype when compared using the Shannon index (Figure 1C, *p*=0.027).

We calculated relative abundance of tumour microbiome and multi-dimensional scaling (MDS) using the Bray-Curtis index to find differences between group (beta-diversity). Unsupervised coordination using PCoA revealed no distinct patterns by PAM50 subtype (Supp Figure 1). We also plotted a Shepherd’s stress plot to measure how well the reduced dimensions reflect the original dissimilarity structure. Relative stress, which is a measure of goodness of fit in MDS and preferably lower value, was 0.29 (Supp Figure 2), indicating that the MDS was a decent fit to the original dissimilarity structure.

In order to examine dissimilarity between groups (inter-group diversity) and the variance contributed by each covariate, we conducted a PERMANOVA analysis, which revealed significant differences between individuals with high versus low IFNγ immune scores (*F*-statistic=2.431, *p*=0.015, Table 1) (Ayers et al. 2017). Dissimilarity contributed by immune scores remained significant when substituted by other immune scores such as Bindea (Bindea et al. 2013) and ESTIMATE (Yoshihara et al. 2013). We also calculated homogeneity using PERMDISP to ensure that differences in group was due to variance and not sample dispersion. All variables examined had homogenous dispersion, with the exception of age at diagnosis (*F*-statistic=1.715, *p*=0.003, Supp. Table 1). It is interesting to note that age at diagnosis explained 19% of variance observed for dispersion. This is expected because patients in this cohort range from 22 – 85 years old, thus resulting in high dispersion.

**Table 1.** PERMANOVA results.

|  | **df** | **SS** | **F** | **Pr(>F)** | **Total**<br>**variance** | **Explained**<br>**variance** |
| --- | --- | --- | --- | --- | --- | --- |
| Model | 12 | 0.87 | 1.03 | 0.391 | 27.69 | 0.031 |
| PAM50 subtype | 4 | 0.31 | 1.12 | 0.283 | 27.69 | 0.011 |
| Age at diagnosis | 1 | 0.085 | 1.21 | 0.227 | 27.69 | 0.0031 |
| Ethnicity | 3 | 0.14 | 0.68 | 0.891 | 27.69 | 0.0052 |
| Stage | 1 | 0.048 | 0.69 | 0.752 | 27.69 | 0.0017 |
| Grade | 1 | 0.078 | 1.11 | 0.323 | 27.69 | 0.0028 |
| Chemotherapy | 1 | 0.060 | 0.86 | 0.535 | 27.69 | 0.0022 |
| IFNγ group | 1 | 0.17 | 2.43 | 0.015 | 27.69 | 0.0062 |
| Residual | 382 | 26.83 | NA | NA | 27.69 | 0.97 |

Inter-group diversity was visualized using supervised ordination with distance-based Redundancy Analysis (dbRDA) which reflected similar findings to PERMANOVA (Figure 2). The figure shows that two axes chosen, RDA1 and RDA2 explained the highest tumour microbiome variance in a multi-dimensional data at 30.9% and 25.2%. The IFNγ immune score had the most significant effect on the variance observed, as confirmed in PERMANOVA.

**Figure 2.**
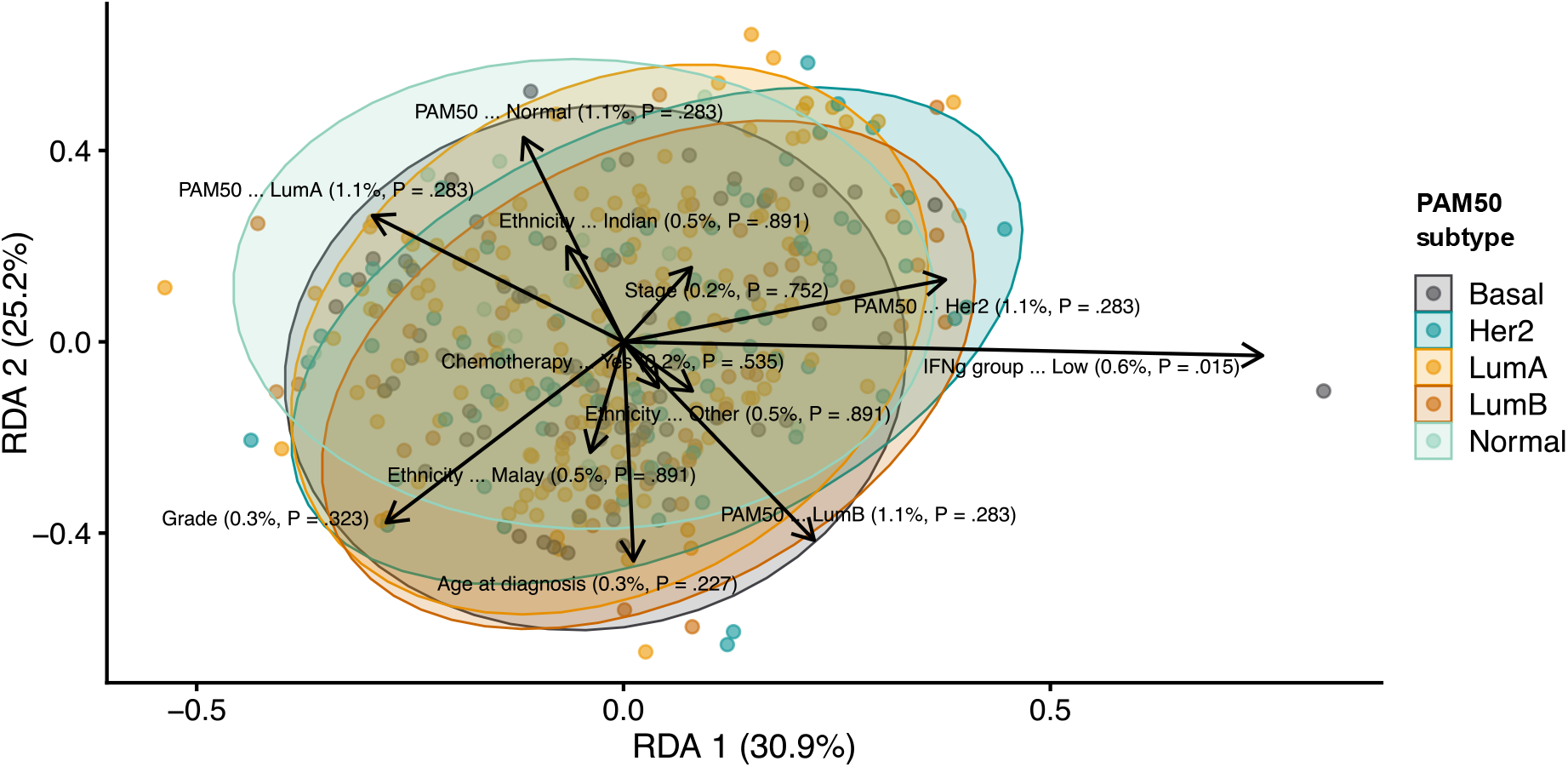
Inter-group diversity of detected intratumoral microbiota, as visualized using supervised ordination with distance-based Redundancy Analysis (dbRDA) against clinical and molecular variables. Clinical and molecular variables included in the analysis were PAM50 molecular subtype, IFN-γ scores (grouped according to median value), ethnicity, chemotherapy treatment (received or not), age at diagnosis, cancer stage, and tumour grade.

### 3.2 Differential abundance analyses of immune scores

Given the previous observation that immune scores had the most significant association with the variance observed in microbial abundance, we investigated which bacteria may be associated with differences in immune scores using differential abundance analysis. Immune scores included in the analysis were Bindea, ESTIMATE, IMPRES, CD8, and IFNγ immune scores as scored in Pan et al. (2020). Immune scores were grouped into high and low by their median. Multiple algorithms, namely ALDEx2, ANCOM-BC2, MaAslin2, Zicoseq, and LinDA, were utilized to search for a consistent pattern while avoiding algorithmic bias towards the identification of differentially abundant bacteria taxa (Nearing et al. 2022). Significant findings were defined as those genera with FDR-adjusted *p*-value<0.05 by two or more algorithms.

These analyses showed that *Sulfidibacter* was significantly increased in patients across most high immune score groups (Bindea, ESTIMATE, CD8 and IFNγ; *p*-value<0.05, Table 2). Additionally, *Priestia* and *Pseudoalteromonas* were significantly increased in IFNγ high groups, while *Bacillus* was significantly increased in patients categorized into low IMPRES score group across at least two separate algorithms.

**Table 2.**
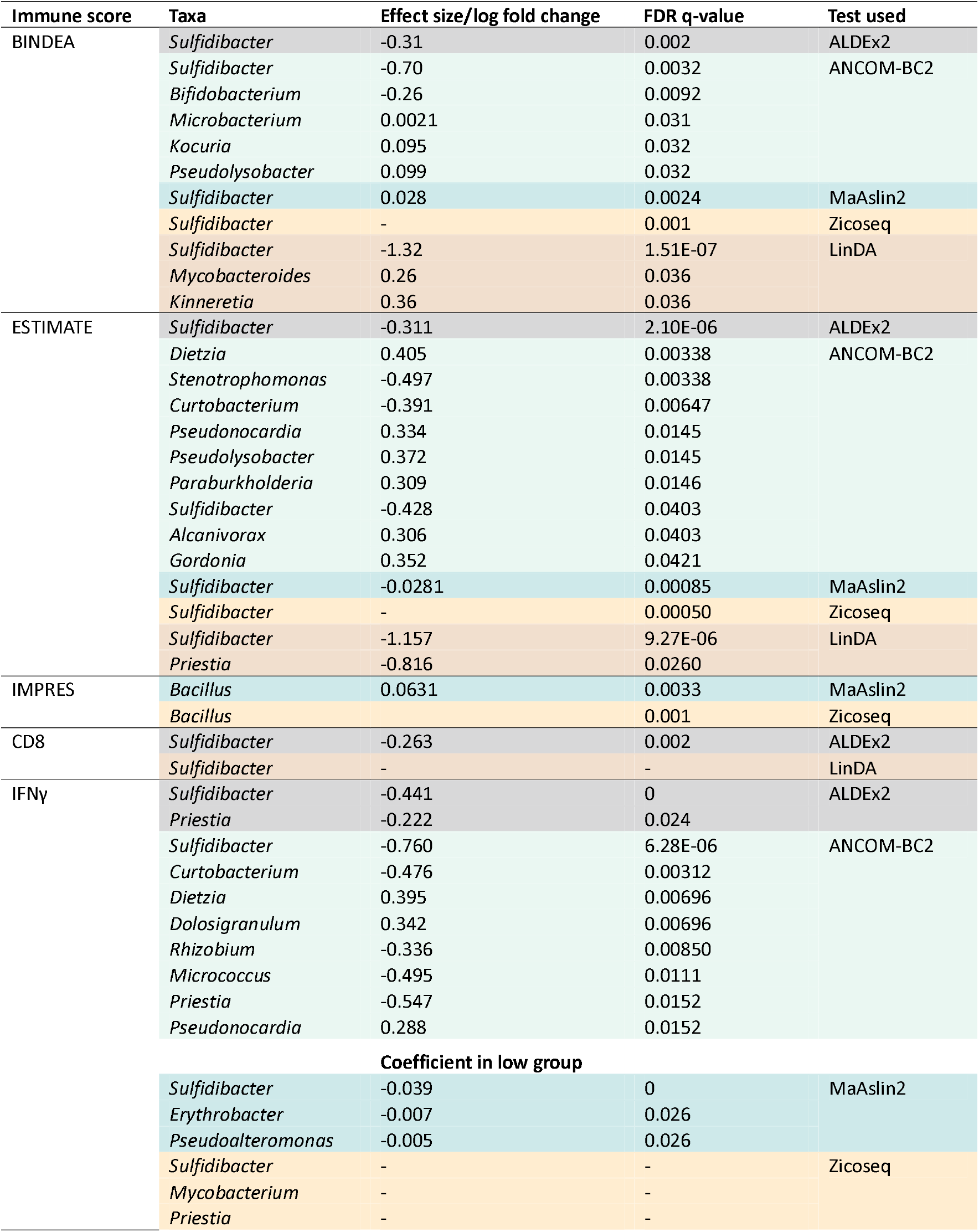
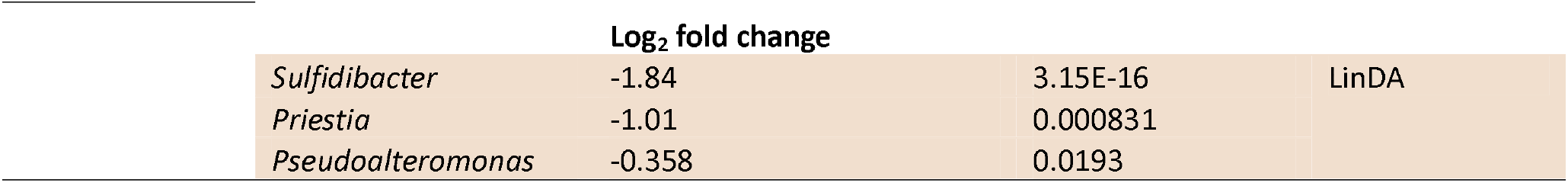
Results for differential abundance analysis for each immune score, the effect size/log fold change and the test used. A negative effect size/log fold change indicates that the taxa was enriched in the group with high immune scores.

### 3.3 Validation of significant associations in a validation cohort

Samples that were sequenced in a later batch were used as a validation cohort (*n*=419). In order to validate our previous finding of an association between the microbial abundance of specific bacterial genera with immune scores, we compared the normalized abundance of *Sulfidibacter, Priestia*, and *Pseudoalteromonas* between samples with high versus low immune scores.

We found that *Sulfidibacter* was significantly higher in abundance among the higher immune score groups for Bindea, ESTIMATE, and IFNγ (Figure 3, *t*-test *p*<0.05), but not CD8 (*p*=0.38), in our validation cohort. Additionally, *Priestia* was also significantly higher in patients with high IFNγ scores in our validation cohort (p<0.0001). However, contrary to our discovery cohort, *Pseudoalteromonas* was not significantly different between IFNγ high and low groups (*p*=0.16). Overall, the results from our validation cohort confirmed most but not all of the associations between bacterial abundance and immune scores from the discovery cohort.

**Figure 3.**
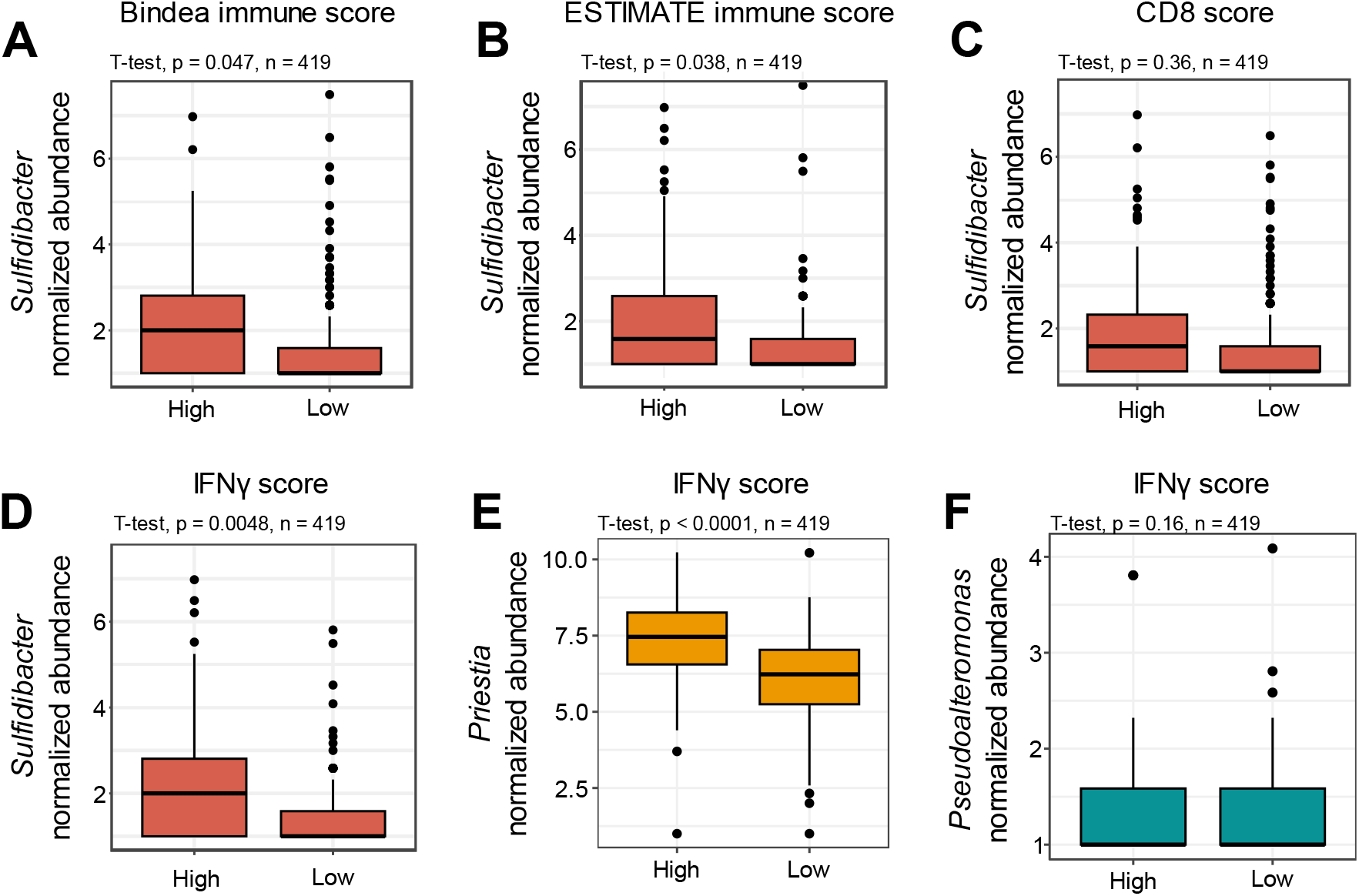
Association of intratumoral microbiota with immune scores in a validation cohort of Asian breast cancer samples (n=419). (A-D) Comparison of RNAseq-derived normalized abundance scores for *Sulfidibacter* between samples with high versus low immune scores according to the median value, for Bindea, ESTIMATE, CD8, and IFN-γ immune scores, respectively. (E-F) Comparison of RNAseq-derived normalized abundance scores for *Priestia* (E) and *Pseudoalteromoas* (F) between samples with high versus low IFN-γ immune scores, according to the median value.

### 3.4 Random Forest prediction of immune scores from microbiome data

We used machine learning to explore the possibility of utilising the tumour microbiota to predict samples with high or low immune scores. A 5-fold cross-validation random forest model was used with an 80-20 split between the training and testing dataset. The random forest model successfully predicted immune high and immune low groups in our full dataset (*n*=883), with an area under the ROC curve (AUC-ROC) of 0.80 for IFNγ, 0.78 for Bindea, 0.72 for ESTIMATE, 0.72 for CD8, and 0.60 for IMPRES (Table 3, Figure 4A). Across all five immune scores analysed, the random forest models of the tumour microbiota were able to predict the immune scores significantly better than chance (AUC-ROC 95% CI > 0.5) and with moderately high sensitivity and specificity in most cases except for IMPRES. The random forest model with the best predictive performance was for IFNγ scores, with an area under the precision-recall curve (AUC-PR) of 0.76 (Figure 4B), and the top three features that contributed to the random forest binary classification model for IFNγ were *Sulfidibacter, Prestia*, and *Erythrobacter* (Figure 4C). Importantly, the tumour microbiome was still significantly predictive for IFNγ scores even when *Sulfidibacter* alone or *Sulfidibacter* and *Priestia* were dropped from the training data (AUROC of 0.72 [95% CI = 0.70-0.76] and 0.70 [95% CI = 0.67-0.73] respectively, Supp. Table 2), suggesting that this association may be robust.

**Table 3.** Random forest prediction metrics for prediction of immune scores using intratumoral microbiome relative abundance scores (n=883).

| Immune score | F1 Score | Area under the ROC curve (95% CI) | Precision (95% CI) | Sensitivity (95% CI) | Specificity (95% CI) |
| --- | --- | --- | --- | --- | --- |
| IFNγ | 0.75 | 0.80 (0.77-0.83) | 0.67 (0.63-0.70) | 0.87 (0.83-0.89) | 0.61 (0.57-0.66) |
| Bindea | 0.72 | 0.78 (0.75-0.81) | 0.70 (0.66-0.74) | 0.74 (0.70-0.78) | 0.70 (0.65-0.73) |
| ESTIMATE | 0.68 | 0.72 (0.69-0.75) | 0.65 (0.61-0.69) | 0.70 (0.66-0.74) | 0.63 (0.59-0.67) |
| CD8 | 0.65 | 0.72 (0.69-0.75) | 0.65 (0.61-0.69) | 0.64 (0.60-0.69) | 0.68 (0.64-0.72) |
| IMPRES | 0.27 | 0.60 (0.54-0.66) | 0.17 (0.14-0.21) | 0.65 (0.56-0.73) | 0.55 (0.51-0.58) |

**Figure 4.**
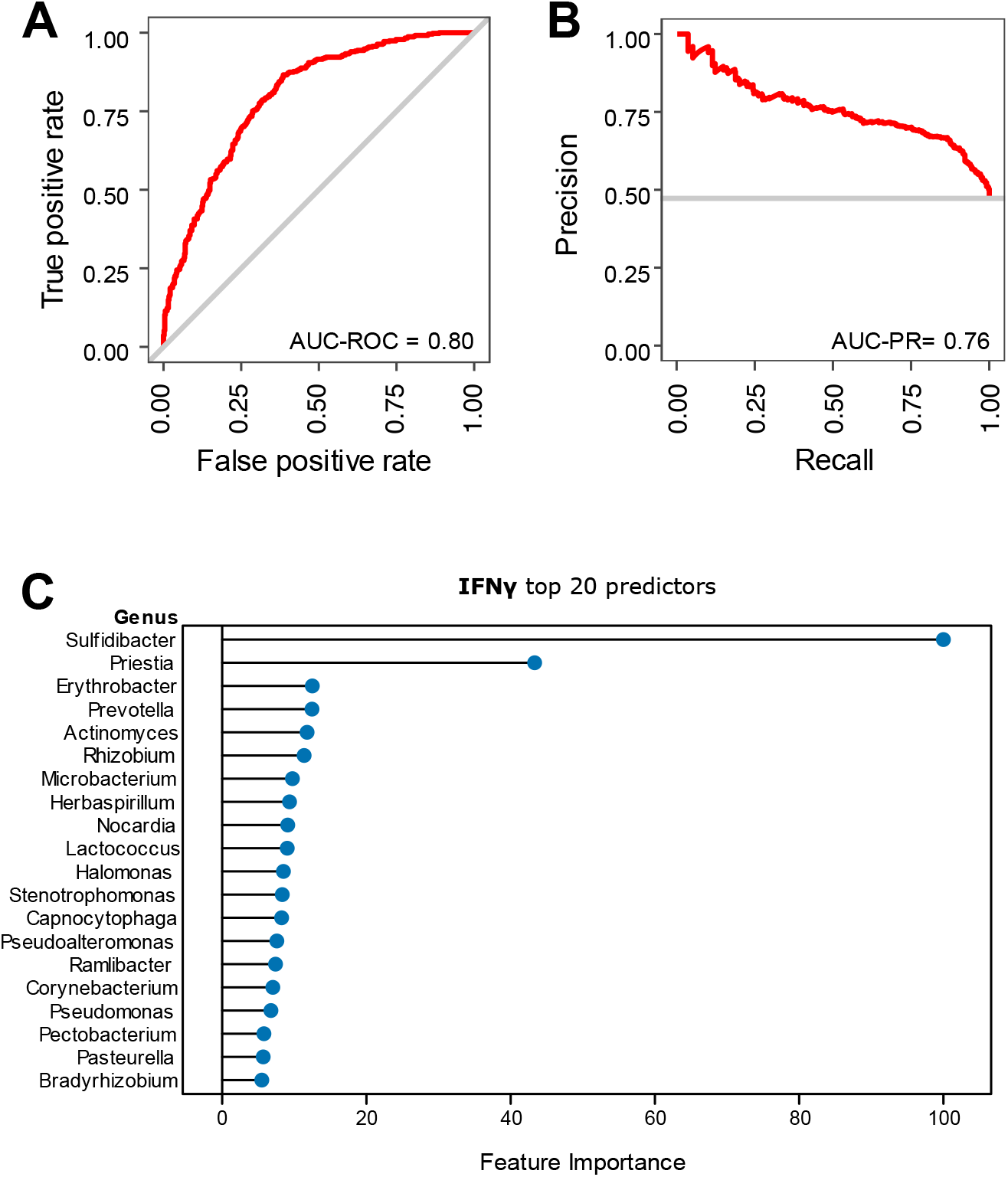
Random forest prediction of IFN-γ scores using intratumoral microbial abundance. Receiver operating characteristics **(A)** and precision-recall curve **(B)** for a random forest model trained to predict IFN-γ scores using intratumoral microbial abundance. **(C)** Top 20 most important features (abundance of specific microbial genera) used by the random forest model to predict IFN-γ scores.

### 3.5 *In vitro* validation

We also sought to validate the existence and overall abundance of the tumour microbiome in our samples using orthogonal methods. Thus, we conducted an *in vitro* validation of microbial abundance using qPCR amplification of the bacterial 16S V3V4 region of 20 randomly-selected samples from our cohort. The estimated total microbial copy numbers derived from qPCR were then compared with microbial read counts derived from RNA sequencing in order to determine their correlation (Figure 5). Both Spearman’s and Pearson’s correlation revealed moderately strong associations between the two variables, with correlation coefficients of 0.513 (*p*=0.020) and 0.7104 (*p*=0.00045) respectively, suggesting that our overall per-sample microbial read counts derived from RNA sequencing were reliable.

**Figure 5.**
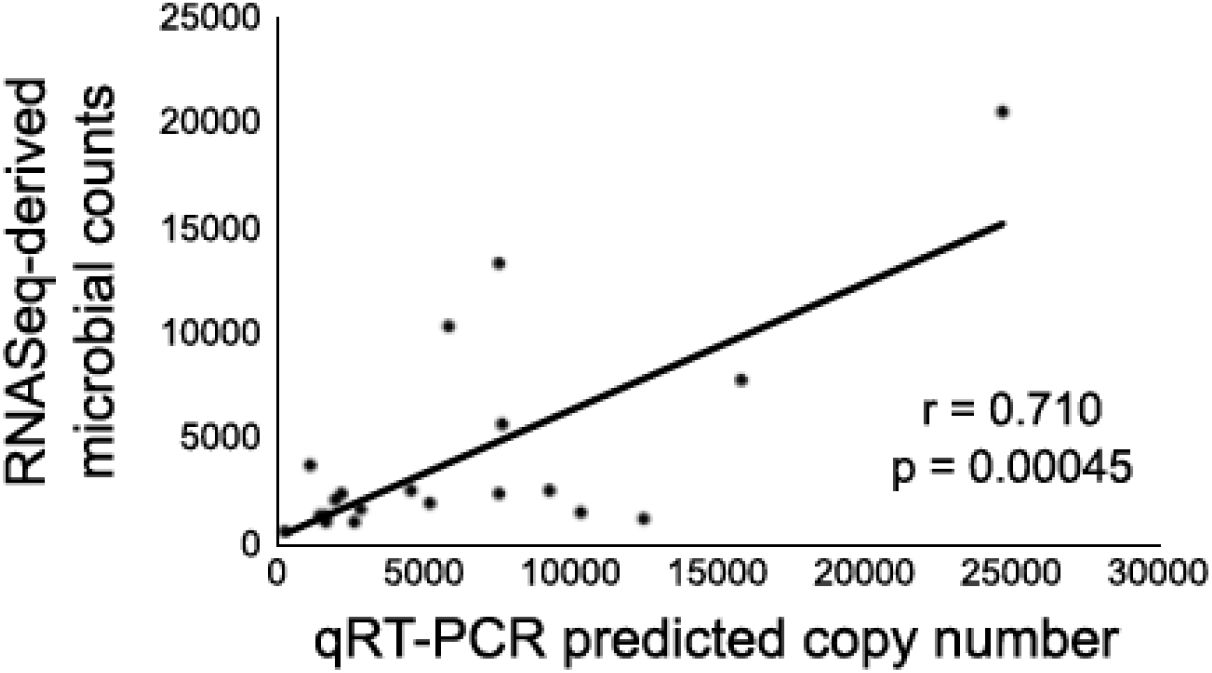
Predicted copy number for the combined intratumoral microbiome, as determined by qRT-PCR amplification of bacterial 16S V3V4, compared to total transcript counts for all detected microbiota derived from RNAseq, for 20 Asian breast cancer tumour samples from the MyBrCa cohort.

## 4.0 Discussion

In this study, we sought to characterize the tumoral microbiota of Asian breast cancer patients to understand its association with molecular subtypes and immune scores. We used a compositional data analysis method involving five algorithms to analyze microbial read counts derived from 558 RNA-seq samples, followed by validation with a separate cohort of 419 samples as well as qPCR of 20 randomly-selected samples. Our findings suggest a lack of association between the intra-tumoral microbiome and most clinical variables, but also suggest a potential association between the intra-tumoral microbiome and immune scores in our Asian cohort.

We observed a largely homogenous intra-group diversity in the Malaysian breast tumor microbiome across PAM50 subtypes, except for the basal subtype which had a significantly more diverse microbiota composition compared to the HER2-enriched subtype. The homogeneity observed is in line with Desalegn et al. (2023) who reported no significant differences in tumour microbiota between PAM50 subtypes among Ethiopian breast cancer patients. Kim et al. (2021) reported similar findings in a Korean cohort, additionally showing two distinct clusters independent of subtypes associated with regional recurrence free survival. Other studies have reported distinct microbiome compositions between tumour and normal adjacent tissue samples (Tzeng et al. 2021, Banerjee et al. 2021, German et al. 2023), which is expected given that comparisons between healthy and diseased microbiomes have consistently reported lower microbial diversity in the latter (Kriss et al. 2018).

However, the significant difference in microbiome diversity between the basal and HER2-enriched subtype has not been reflected in current literature. Interestingly, Chen et al. (2020) reported that Asian breast cancer patients are less likely to have luminal A and basal subtypes but more likely to have luminal B and HER2-enriched subtypes than Western patients. Tumour microbiomes tend to be less well-characterized compared to the gut microbiome. This gap in knowledge is further exacerbated by the lack of Asian-centric cohorts (Li et al. 2022; Luo et al. 2023; Kim et al. 2021). Furthermore, tumour microbiome studies with multiethnic cohorts tend to have a relatively low representation of Asians (German et al. 2023, Parida et al. 2023). Considering the sample size of our cohort, it is possible that the observed differences in microbiome diversity between basal and HER-2 enriched subtypes could be specific to Asian populations but this requires further validation.

The results of our inter-group diversity analysis further revealed that variation in the Malaysian breast tumour microbiome was significantly associated with immune scores, while molecular subtype, age at diagnosis, cancer stage, tumour grade, ethnicity, and treatment type had no association with the variation found in the Malaysian breast tumour microbiome.

There is evidence that microbes can interact with cells patrolling the tumour microenvironment, notably close interactions with immune cells possibly affecting tumour inhibition and proliferation (Ma et al. 2021). Microbes have been found intracellularly in both cancer and immune cells (Nejman et al. 2020). In other cancers such as pancreatic ductal adenocarcinoma, a uniquely diverse composition of tumour microbiome distinct from that of adjacent healthy pancreatic tissue was found to be associated with more sustained CD8^+^ T cell response in the tumor microenvironment (Riquelme et al. 2019).

Increased immune cell response and variations in immune scores have also been attributed to microbe-derived metabolites present in the tumour immune microenvironment (TIME) (Ma et al. 2021). Short chain fatty acids (SCFAs), such as butyric acid, are known microbe-derived metabolites which can accumulate within tumors and inhibit histone deacetylases (HDACs), referring to chromatin regulatory factors expressed abnormally in a variety of human cancers (He et al. 2021). Butyrate-mediated HDAC inhibition causes the upregulation of transcriptional regulator ID2, triggering the IL-12R signalling pathways in CD8^+^ T cells (Luu et al. 2018). This results in an increased CD8^+^ T cell density and activation in the TIME.

In the case of IFNγ immune score, studies have reported that some bacterial genera can promote IFNγ secretion, including a recently defined community of 11 bacteria that induced IFNγ production preferentially in CD8^+^ T cells in the absence of immunotherapy (Tanoue et al. 2019; Wang et al. 2017). IFNγ secretion in the TIME have also been linked to *Bifidobacterium* (Rezasoltani et al. 2020). One such metabolite is inosine, a purine metabolite which induces naïve T cells to differentiate into CD4^+^ Th1, leading to increased CD8^+^ T-cell infiltration and IFNγ secretion, especially in combination with PD-L1 blockade (Mager et al. 2020; He et al. 2021). It is worth noting that while IFNγ is classically associated with anti tumour effects, IFNγ can upregulate proliferative signals and allow tumour cells to escape recognition by immune cells under certain conditions (Zaidi & Merlino 2011).

Our differential abundance analyses using center log-ratio transformed counts and five different algorithms showed that *Sulfidibacter* was significantly increased in patients with higher immune scores, including Bindea, ESTIMATE, and IFNγ. Additionally, *Priestia* was also more abundant in patients with high IFNγ scores in both our discovery and validation cohorts.

The presence of *Sulfidibacter* in the tumour microbiome was unexpected as it is a novel marine bacterium first isolated and identified from corals (Wang et al. 2022). To date, Wang et al. (2022) is the only publication available which characterizes *Sulfidibacter*. However, given it was proposed as a species of Acidobacteria and members of this phylum are typically associated with aquatic, terrestrial, and extreme environments, *Sulfidibacter* is not expected nor likely to appear in human species. Hence, it is possible that its presence in our data is the result of taxonomic misclassification due to database contamination by human reads or other contamination instead of a true biological signal (Gihawi et al. 2023).

*Priestia* is another marine bacterium previously reported as an arginase producer, an enzyme with potential in cancer treatment by arginine deprivation therapy (Jiao et al. 2023). *Priestia* was previously identified in a Slovakian breast tumour cohort by Hadzega et al. (2021), who conducted transcriptomic sequencing to investigate the breast tumour microbiome and found that *Priestia* was enriched in breast tumours from patients compared to normal tissues from cancer-free women.

Moving forward, our results may have implications for future treatment strategies to modulate IFNγ in the TIME via manipulation of the tumour microbiome. Already, engineered bacteria injected at tumour sites have been found to trigger IFNγ expression through a cascade of pathways that increases anti-tumour effects (Chen et al. 2022). Similarly, Kim et al. (2017) demonstrated the use of gram-negative bacteria outer membrane vesicles to induce anti-tumour effects through the production of anti-tumour cytokines such as IFNγ and CXCL10.

One of the strengths of our study, aside from the sizeable cohort, is the analysis strategy used to mediate batch effect. Batch effect has been and will continue to be a major issue with the rise of big data and large microbiome cohorts. Several strategies to correct it have been reported in literature including conditional quantile regression (Ling et al. 2022), MBECS (Olbrich et al. 2023), Limma, and ComBat. Still, there persists the question of whether these strategies could overcorrect data to the point of distorting data dispersion, resulting in the detection of false positive signals or the masking of true positive signals (Sepich-Poore et al. 2024). To avoid such data distortion, we adapted a strategy from Sepich-Poore et al. (2024) where we used one sequencing batch as an exploratory cohort and another batch as an independent validation cohort.

Our study does have some limitations. The dataset was not initially designed for microbiome investigations, and thus, there is a lack of microbiome controls to rule out environmental contamination. We attempted to reduce the effect of this on our findings by applying appropriate prevalence filtering, testing differential association on five different algorithms, and incorporating the use of a sizable validation cohort. We have also conducted orthogonal validation via qPCR on tumour DNA. However, we cannot completely rule out the presence of contaminants or false positives in our data. Additionally, the inclusion of other omics such as metabolomics, genomics, and gut metagenomics could provide more insights into understanding of the human microbiome and its role in association with cancer.

## Acknowledgements

Cancer Research Malaysia receives charitable funding from the Khind Starfish Foundation, the Ong Hin Tiang & Ong Sek Pek Foundation, Yayasan PETRONAS, and Yayasan Sime Darby which contributed to the funding of this study.

## Author Contributions

LFY contributed to study design, data collection and processing, and drafting of the manuscript and figures. AWYL and JSFC contributed to the drafting of the manuscript and literature review. PYET contributed to data collection, data analysis, and drafting of the manuscript and figures. BKBL contributed to study conceptualization and design as well as data collection. JL contributed to study design and project supervision. SHT also contributed to study design and project funding, as well as project direction and supervision. JWP contributed to study design, project direction and supervision, data analysis and drafting of the manuscript and figures. The work reported in the paper has been performed by the authors, unless clearly specified in the text.

