## Supplementary material for "Detection of intra-tumoral microbiota from transcriptomic sequencing of Asian breast cancer": Supp. Table 1

Supp Table 1. PERMDISP results indicating that PERMANOVA significance is from variance in microbiota and not from dispersion.

|  | **df** | **SS** | **MS** | **F** | **N. Perm** | **Pr(>F)** | **Total variance** | **Explained variance** |
| --- | --- | --- | --- | --- | --- | --- | --- | --- |
| PAM50 | 4 | 0.0063882 | 0.0015971 | 0.2266889 | 999 | 0.934 | 3.141474 | 0.0020335 |
| Age at diagnosis | 54 | 0.6620194 | 0.0122596 | 1.7148757 | 999 | 0.003 | 3.478719 | 0.1903055 |
| Ethnicity | 3 | 0.0232414 | 0.0077471 | 1.1191940 | 999 | 0.347 | 3.075868 | 0.0075560 |
| Stage | 4 | 0.0432895 | 0.0108224 | 1.5720610 | 999 | 0.198 | 3.017259 | 0.0143473 |
| Grade | 2 | 0.0029800 | 0.0014900 | 0.2103359 | 999 | 0.808 | 2.857794 | 0.0010428 |
| Chemotherapy | 1 | 0.0000582 | 0.0000582 | 0.0081558 | 999 | 0.919 | 3.010703 | 0.0000193 |
| IFNγ group | 1 | 0.0002145 | 0.0002145 | 0.0312037 | 999 | 0.884 | 3.079575 | 0.0000696 |
