## Supplementary material for "Detection of intra-tumoral microbiota from transcriptomic sequencing of Asian breast cancer": Supp. Table 2

Supp. Table 2. Random forest prediction metrics for prediction of IFNγ scores using intratumoral microbiome relative abundance scores (n=883) when *Sulfidibacter* (first row) or *Sulfidibacter* and *Priestia* (second row) were excluded from the training data.

| **Condition** | **F1 Score** | **Area under the ROC curve**  **(95% CI)** | **Precision**  **(95% CI)** | **Sensitivity**  **(95% CI)** | **Specificity**  **(95% CI)** |
| --- | --- | --- | --- | --- | --- |
| *Sulfidibacter* excluded | 0.66 | 0.73 (0.70-0.76) | 0.68 (0.64-0.73) | 0.64 (0.59-0.68) | 0.74 (0.70-0.77) |
| *Sulfidibacter* and *Priestia* excluded | 0.62 | 0.70 (0.67-0.73) | 0.64 (0.59-0.68) | 0.60 (0.56-0.65) | 0.69 (0.65-0.73) |
