## Supplementary material for "Detection of intra-tumoral microbiota from transcriptomic sequencing of Asian breast cancer": Supp. Figure 1, Supp. Figure 2

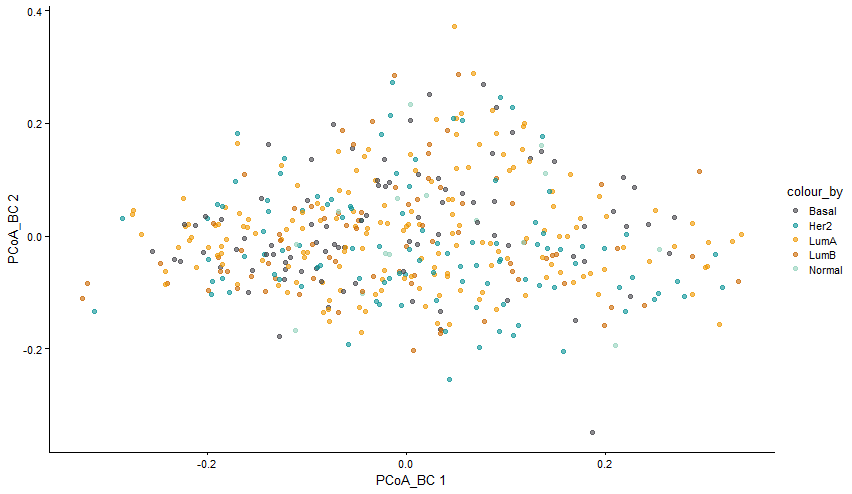

**Supp Figure 1.** Unsupervised coordination using principal component analysis of the MyBrCa tumor microbiome relative abundance scores by genera, colour coded by PAM50 subtype.

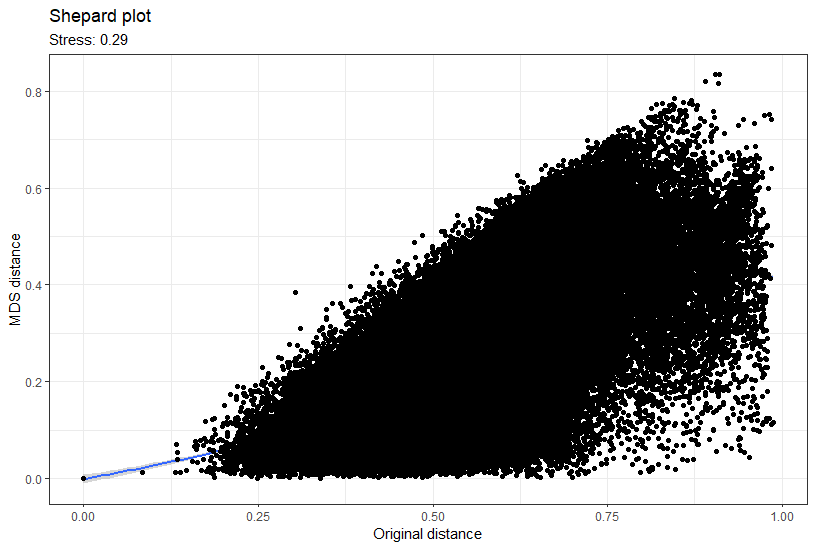

**Supp Figure 2**. Shepard’s stress test plot for the MyBrCa tumor microbiome relative abundance scores by genera. Relative stress was 0.29.
